# Modulation of endothelial organelle size as an antithrombotic strategy

**DOI:** 10.1101/2020.05.16.099580

**Authors:** Francesco Ferraro, Joana R. Costa, Robin Ketteler, Janos Kriston-Vizi, Daniel F. Cutler

## Abstract

It is long-established that Von Willebrand Factor (VWF) is central to haemostasis and thrombosis. Endothelial VWF is stored in cell-specific secretory granules, Weibel-Palade bodies (WPBs), uniquely rod-like exocytic organelles generated in a wide range of lengths (0.5 to 5.0 µm). It has been shown that WPB size responds to physiological cues and pharmacological treatment and that, under flow, VWF secretion from shortened WPBs produces a dramatic reduction of platelet and plasma VWF adhesion to an endothelial surface. WPB-shortening therefore represents a novel target for antithrombotic therapy acting via modulation of VWF adhesive activity. To this aim, we screened a library of licenced drugs and identified several that prompt WPB size reduction. These compounds therefore constitute a novel set of potentially antithrombotic compounds.

**Summary:** The size of the endothelial secretory granules that store Von Willebrand Factor correlates with its activity, central to haemostasis and thrombosis. Here, human-licenced drugs that reduce the size of these secretory granules are identified, providing a set of novel potential anti-thrombotic compounds.

## Introduction

Endothelial Von Willebrand Factor (VWF) plays a fundamental role in haemostasis, with deficiencies in its activity causing von Willebrand Disease (VWD), the most common inherited human bleeding disorder (Sadler, 1998). VWF is a large multi-domain glycoprotein, whose function in haemostasis depends on its multimeric status. VWF multimers act as mechano-transducers, which respond to shear forces in the circulation by stretching open and exposing binding sites for integrins, collagen, platelets and homotypic interaction (i.e., between VWF multimers) (Ruggeri and Mendolicchio, 2015). Endothelial cells secrete VWF in a highly multimerized form, known as ultra-large (UL)-VWF, highly sensitive to haemodynamic forces and thus very active in platelet binding (Zhang et al., 2009).

UL-VWF’s potential to cause spontaneous thrombus formation is controlled by a circulating protease, ADAMTS13, which generates the less-multimerised, less active forms of VWF seen in plasma (Zhang et al., 2009). Persistence of UL-VWF in the circulation leads to microvascular thrombosis and the highly morbid and potentially life threatening clinical manifestations observed in a host of infectious and non-infectious diseases, such as sepsis and thrombotic thrombocytopenic purpura (TTP)(Chang, 2019; Tsai, 2010). Although these may be the most extreme examples of excess VWF function, many common disorders including hypertension and diabetes are characterised by increased VWF plasma levels (Apostolova et al., 2018; Westein et al., 2017).

While VWF is stored in the secretory granules of both platelets and endothelial cells, most of the VWF circulating in plasma derives from endothelial WPBs and is fundamental to haemostasis (Kanaji et al., 2012). Vessel injury, in either the macro- or microcirculation, triggers localized stimulated exocytosis of WPBs, mediated by a variety of agonists (McCormack et al., 2017). UL-VWF secreted from activated endothelium forms cable-like structures built of multiple multimers, both in vitro and in vivo (Arya et al., 2002; Rybaltowski et al., 2011). These VWF “strings” provide scaffolds for the recruitment of circulating platelets and soluble plasma VWF, contributing to the formation of the primary haemostatic plug, but also potentially promoting microangiopathy (Nicolay et al., 2018; Ruggeri and Mendolicchio, 2015).

The length of VWF strings generated upon exocytosis reflects the size of the WPBs in which VWF was stored. WPBs are cigar-shaped organelles, whose length ranges ten-fold, between 0.5 and 5.0 µm. Their size depends on the structural status of the endothelial Golgi apparatus where they form, and experimental manipulations causing Golgi fragmentation consistently result in the formation of only short WPBs (Ferraro et al., 2014). Short WPBs also form when VWF biosynthesis by endothelial cells is reduced, or following statin treatment, via Golgi fragmentation-independent and –dependent mechanisms, respectively (Ferraro et al., 2014; Ferraro et al., 2016). The metabolic status of endothelial cells also regulates WPB size through an AMPK-mediated signalling pathway (Lopes-da-Silva et al., 2019).

Importantly, *in vitro* experiments have revealed that reducing WPB size results in the shortening of the VWF strings they generate and in much-reduced recruitment of platelets and soluble circulating VWF to the endothelial surface (Ferraro et al., 2014; Ferraro et al., 2016). Conversely, endothelial cells respond to raised glucose levels (mimicking hyperglycemia) by producing longer WPBs, suggesting a link between long VWF strings and thrombotic manifestations in diabetes (Lopes-da-Silva et al., 2019), which is often associated with high levels of plasma VWF and microangiopathy (Westein et al., 2017).

WPB size is, therefore, plastic and responds to physiological cues and pharmacological treatments. Such findings suggest that drug-mediated reduction of WPB size might provide alternative or coadjuvant therapeutic approaches to current clinical interventions in thrombotic pathologies where dysregulated formation and/or prolonged persistence of VWF strings on vascular walls play a triggering role.

We designed a screen to identify drugs that reduce WPB size and thus can potentially reduce endothelial pro-thrombotic capacity. Out of 1280 human licensed drugs we found 58 compounds fitting our criteria, with a variety of mechanisms of action consistent suggesting a number of pathways that influence biogenesis of WPBs.

## Results

### Screen

A quantitative high-throughput microscopy-based workflow, dubbed high-throughput morphometry (HTM), allows rapid quantification of the size of tens to hundreds of thousands of WPBs within thousands of endothelial cells (Ferraro et al., 2014). HTM has been applied for analytical purposes and in phenotypic screens (Ferraro et al., 2014; Ferraro et al., 2016; Ketteler et al., 2017; Lopes-da-Silva et al., 2019; Stevenson et al., 2014). In the present report, HTM was deployed to identify compounds that can induce a reduction of WPB size in Human Umbilical Vein Endothelial Cells (HUVECs). WPBs longer than 2 µm represent ∼ 20% of these VWF-storing organelles, but contain roughly 40% of all endothelial VWF. Thus, while a minority, these long WPBs are disproportionally important with respect to secreted endothelial VWF (Ferraro et al., 2014). For the purpose of the screen, we quantified WPB size as the ratio between the area covered by WPBs longer than 2 µm and the area covered by all these organelles; we define this parameter as “fractional area (FA) of long WPBs” (**Figure 1A;** (Ferraro et al., 2016). The effect of each compound in the library on WPB size was compared to two controls: treatment with DMSO and Nocodazole, the negative and positive controls, respectively, for WPB shortening (Ferraro et al., 2014; Ferraro et al., 2016) (**Figure 1B and 1C**). HUVECs were incubated with the 1280 compounds of the Prestwick library at 10 µM for 24 h in single wells of 96-well plates, in duplicate (two separate plates; **Figure 1D**). After fixation, immuno-staining and image acquisition, the FA of long WPBs in treated cells was quantified. Per plate and inter-plate normalization to DMSO-controls was implemented in order to rank the effects of the library compounds on WPB size by Z-score, (**Figure 1E**).

**Figure 1.**
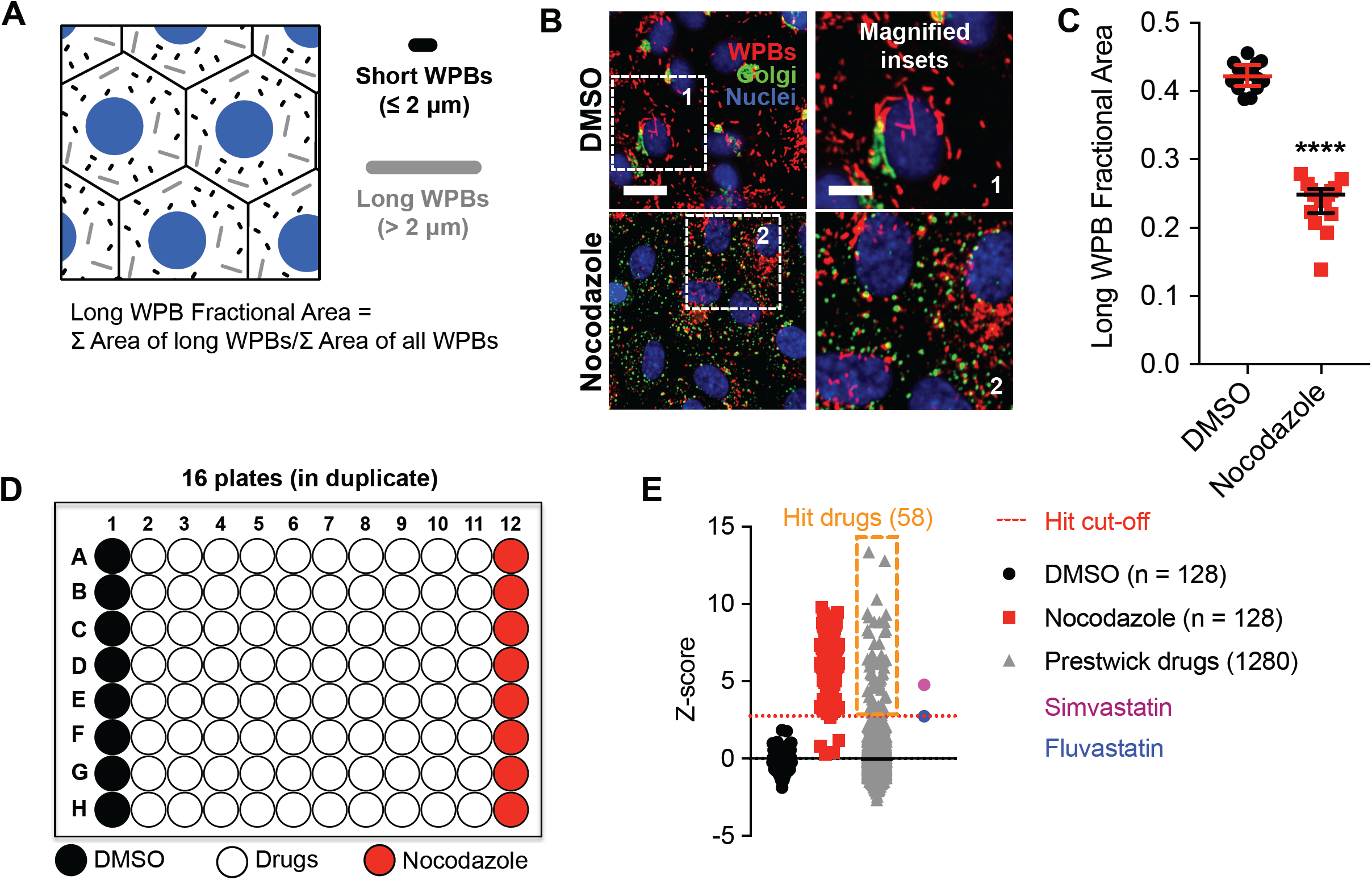
Drug screen: conception, execution and results. **A**. WPB sizes range between 0.5 and 5 µm. To measure a single quantitative parameter accounting for WPB size in a organelle population, we calculated the “fraction of the area (FA) covered by long (i.e., > 2 µm) WPBs. **B**. DMSO and Nocodazole were used as negative and positive control treatments, respectively, for reducing WPB size. Micrographs show the effects of the two control treatments on HUVECs (24 h); scale bar: 20 µm (inset, 10 µm). **C**. Quantification of the “FA of long WPBs” for cells treated as in B. Data-points represent the values calculated for each well of a 96-well plate; median and interquartile ranges are shown. ****, P < 0.0001; Mann-Whitney test. **D**. Screen setup. Individual library drugs were dispensed into single wells of 96-well plates, which also had two columns treated with DMSO and Nocodazole controls. Two plates with identical drug layout (biological duplicates) were analyzed. **E**. Screen results. Z-scores of the library drugs are plotted. Hit drugs were selected based on the effects of fluvastatin and simvastatin, which we previously showed to reduce WPB size. Fluvastatin, with the lowest Z-score, was used as cut-off for the selection of hit compounds.

### Hit selection

Statins are cholesterol-lowering drugs that inhibit the enzyme 3-hydroxy, 3-methylglutaryl CoA reductase (HMGCR) in the mevalonate pathway, upstream of the biosynthesis of cholesterol. We have previously shown that treatment of endothelial cells with two statins, simvastatin and fluvastatin, induces WPB size shortening, resulting in reduced adhesive properties of the VWF released by activated endothelial cells (HUVEC), measured by the reduced size of platelet-decorated VWF strings and by the recruitment of VWF from a flowing plasma pool (Ferraro et al., 2016). Fluvastatin and simvastatin are present in the Prestwick library. We therefore used their Z-score to establish a stringent cut-off for selection of positive hits. This approach identified 58 compounds, 4.5% of the library (**Figure 1E**, orange box, simvastatin and fluvastatin). Of note, aside from simvastatin and fluvastatin, the other three statins present in the Prestwick library were among the selected hits (**Table 1**), indicating that both our screening approach and the criterion for hit selection are robust.

**Table 1.** Licenced drugs with WPB-shortening activity. Individual drugs were assigned to pharmacological classes, indicated by numbers, based on their mechanism of action (see Appendix Table). 1, cardiac glycoside or cardenolide (Na^+^/K^+^-ATPase inhibitor); 2, statin (3-hydroxy, 3-methylglutaryl CoA reductase, HMGCR inhibitor; 3, topoisomerase inhibitor/ DNA-intercalating agent; 4, serotonin (5-HT) receptor antagonist; 5, histone deacetylase (HDAC) inhibitor; 6, dopamine receptor antagonist; 7, histamine receptor antagonist; 8, microtubule depolymerizing compound; 9, pH gradient-depleting compound; 10, multidrug resistance protein 1 (MRP1/ABCB1) inhibitor.

|  |  |  |  |  |  |
| --- | --- | --- | --- | --- | --- |
| Digoxin | 1 | Thiopropazine | 4, 6 | Carvedilol | 10 |
| Lanatoside C | 1 | Fluspirilene | 4, 6 | Hexachlorophene |  |
| Digoxigenin | 1 | Fluphenazine | 4, 6, 10 | Clomiphene |  |
| Proscillaridin A | 1 | Clomipramine | 4, 10 | Alclometasone |  |
| Digitoxigenin | 1 | Sertindole | 4, 6 | Ciclesonide |  |
| Lovastatin | 2, 5, 10 | Clemizole | 4, 7 | Tegaserod |  |
| Atorvastatin | 2, 5, 10 | Trifluoperazine | 6, 10 | Aprepitant |  |
| Simvastatin | 2, 5, 10 | Prochlorperazine | 6, 10 | Chlormezanone |  |
| Fluvastatin | 2, 5 | Astemizole | 7, 10 | GBR 12909 |  |
| Mevastatin | 2 | Vorinostat | 5 | Cycloheximide |  |
| Mitoxantrone | 3, 10 | Colchicine | 8, 10 | Adenosine 5'-monophosphate |  |
| Camptothecine | 3, 10 | Nocodazole | 8 | Methyldopa (L,-) |  |
| Etoposide | 3, 10 | Monensin | 9 | Perhexiline |  |
| Daunorubicin | 3, 10 | Ambroxol | 9 | Parbendazole |  |
| Doxorubicin | 3, 10 | Thonzonium | 9 | Pyrvinium |  |
| Epirubicin | 3 | Bepiridil | 10 | Loperamide |  |
| Podophyllotoxin | 3, 8 | Cyclosporin A | 10 | Dilazep |  |
| Clofazimine | 3, 10 | Itraconazole | 10 | Verteporfin |  |
| Quinacrine | 3, 10 | Imatinib | 10 |  |  |
| Maprotiline | 4, 6, 7, 10 | Azithromycin | 10 |  |  |

### Hit pharmacology

Forty-one (71%) of the drugs identified could be allocated in ten pharmacological classes, each including at least three compounds (**Table 1; and Supplemental Table 1**), consistent with a variety of mechanisms of WPB size control. Some of the compounds are shared by more than one class (**Table 1 and Figure 2A**); an indication that, aside from their known molecular targets, these drugs may exert their effect on WPB size through a yet unidentified common cellular pathway.

**Figure 2.**
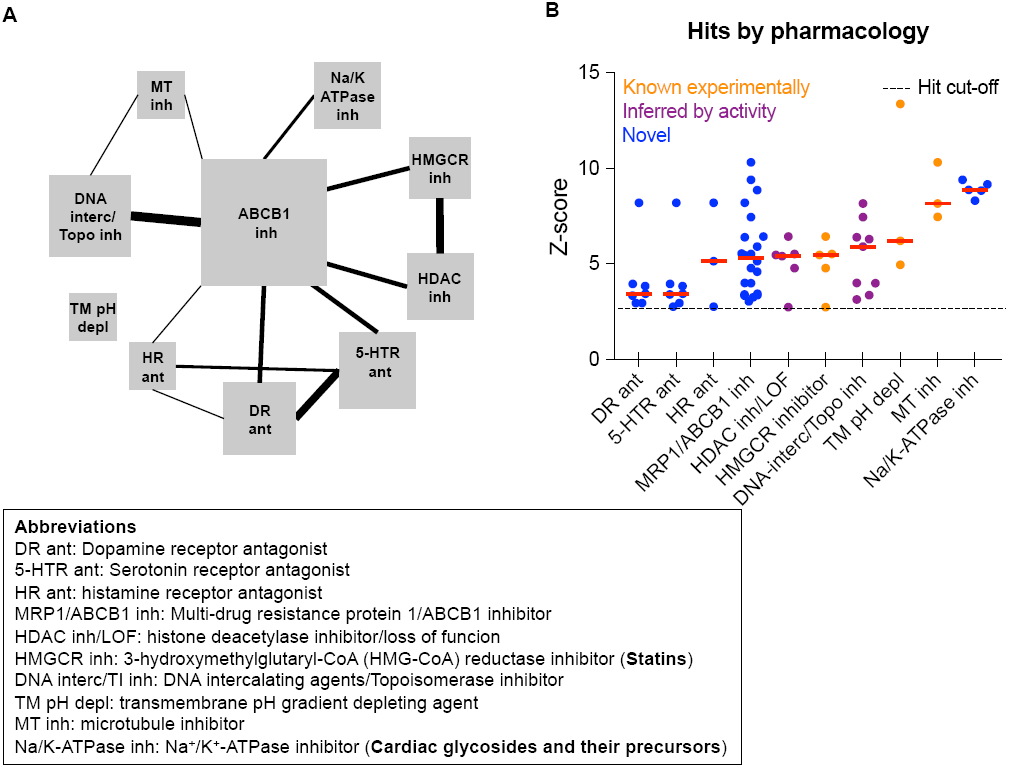
WPB size-reducing drugs and their potency as pharmacological classes. **A.** Graphical summary of the known mechanism of action (MoA) of the hits reported in the Supplemental Table 1. Each pharmacological class is depicted by a square, whose area is proportional to the number of compounds it includes. Lines connecting the squares represent common drugs between pharmacological classes. Thickness of the lines indicates the number of drugs shared. **B**. Drug classes ranked by potency, using their median Z-score in the screen. Each data-point represents one drug and classes are color-labeled based on their know, inferred from the literature and previously unknown (novel) effects on WPB size.

Among the pharmacological classes capable of inducing WPB shortening, we found microtubule (MT) depolymerizing agents, histone deacetylase (HDAC) inhibitors, topoisomerase inhibitors and, as mentioned earlier, statins (HMGCR inhibitors). Compounds with these mechanisms of action have been shown induce unlinking of the ribbon architecture of the Golgi apparatus, i.e. Golgi “fragmentation” (Farber-Katz et al., 2014; Ferraro et al., 2016; Gendarme et al., 2017; Thyberg and Moskalewski, 1985). Since an intact Golgi ribbon is required for the biogenesis of long WPBs (Ferraro et al., 2014) (see nocodazole in **Figure 1B**), identification of these classes of molecules in our screen was expected. Work from our lab also showed that neutralization of the acidic lumen of WPBs disrupts the tubular structure of VWF, shifting the organelle shape from cylindrical to spherical, therefore detected as shortening (Michaux et al., 2006); and, indeed, we identified transmembrane pH gradient depleting agents. Reduction in VWF biosynthesis results in shorter WPBs without affecting the Golgi architecture (Ferraro et al., 2014) and one of the screen hits, cycloheximide, is a classic protein synthesis inhibitor.

Aside from those expected, entirely novel WPB-shortening drug classes were also identified, including neurotransmitter receptor antagonists and cardiac glycosides. Interestingly, several compounds, beside their known mechanism of action, also inhibit multidrug resistance protein 1, MRP1/ABCB1 (**Table 1, Figure 2A and Supplemental Table 1**), which may hint at the common cellular pathway discussed above. The cardiac glycosides and their cardenolide precursors were prominent among these novel pharmacological classes. These molecules, which inhibit Na^+^/K^+^-ATPase (**Table 1** and **Figure 2A**), have a long clinical history in the treatment of congestive heart failure and atrial fibrillation and have recently attracted interest as potential anticancer molecules (Newman et al., 2008) and senolytics, i.e., selective inducers of senescent cell death (Triana-Martinez et al., 2019). Cardiac glycosides and cardenolides display a powerful WPB shortening effect and induce Golgi apparatus compaction (**Figures 2B and 3A**) instead of its fragmentation, suggestive of a novel WPB size-reducing mechanism. In most cases, sub-micromolar concentrations of these compounds were sufficient to reduce WPB size (**Figure 3B**).

**Figure 3.**
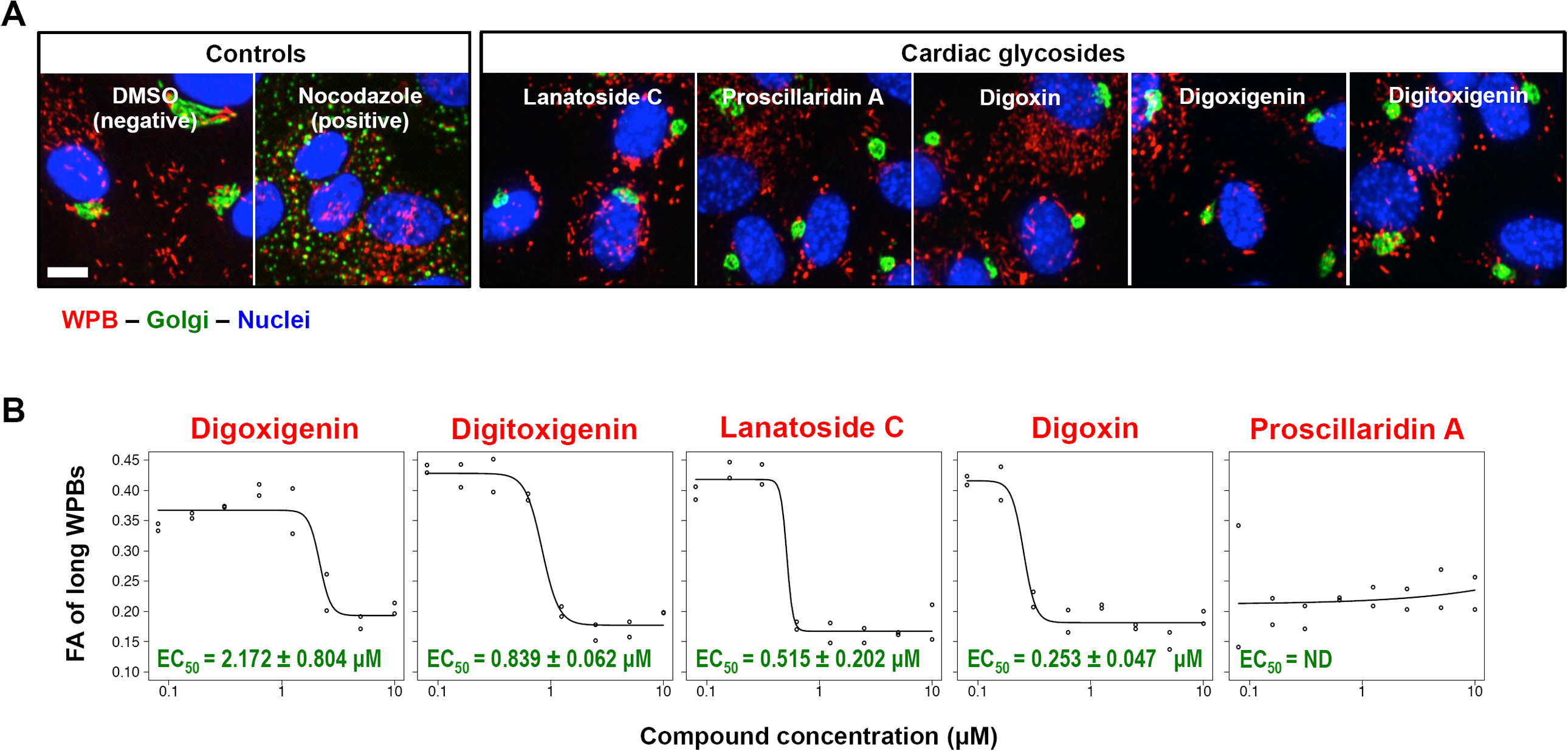
Effects of cardiac glycosides and cardenolides. **A**. HUVECs were treated for 24 h with 10 µM each of the indicated compounds; original micrographs from the screen are shown. These compounds induce WPB-shortening and Golgi apparatus compaction (compare to DMSO negative control). Scale bar: 10 µm. **B**. The effects of cardiac glycosides and cardenolides on WPB size were measured for a range of concentrations (10 µM to 78 nM; two-fold dilutions); calculated EC_50_ values are reported (from highest to lowest; right to left). WPB size remained small in HUVECs treated with proscillaridin A at all the concentrations tested and its EC_50_ could not be determined (ND), indicating that it is likely to be found at low nM or pM concentrations.

## Discussion

Upon exocytosis, endothelial UL-VWF self-assembles into strings, which serve as recruiting platform for platelets and circulating VWF, thus promoting the formation of the primary haemostatic plug (Ferraro et al., 2016; Ruggeri, 2007; Varga-Szabo, 2008). VWF-strings also mediate pathological processes such as tumour metastasis, endocarditis and microangiopathies (Bauer et al., 2015; Nicolay et al., 2018; Pappelbaum et al., 2013). Interventions reducing the persistence and/or activity of VWF-strings are therefore of interests as potential anti-thrombotic therapies.

Modulation of organelle size has been suggested as a potential strategy to regulate biological functions and correct pathological states (Marshall, 2012). In this context, WPBs represent a paradigmatic example. Reduction of WPB size has no effect on UL-VWF formation, but blunts generation of long platelet-decorated VWF-strings following exocytosis and recruitment of plasma VWF to the endothelial surface (Ferraro et al., 2014; Ferraro et al., 2016). Interventions that shorten WPBs could therefore provide alternative or coadjuvant therapies to clinical interventions in thrombotic pathologies where dysregulated formation and/or prolonged persistence of VWF-strings play a triggering role.

Apart from pharmacological treatments and other experimental manipulations disrupting the integrity of the Golgi apparatus (Ferraro et al., 2014; Ferraro et al., 2016), formation of short WPBs is mediated by endogenous signalling pathways. We have uncovered a pathway involving AMPK-dependent regulation of the Arf-GEF GBF1, which is independent of alterations in the structure of the Golgi apparatus and links WPB size to the metabolic status of endothelial cells (Lopes-da-Silva et al., 2019). Small WPBs are also generated upon overexpression of the transcription factor KLF2 (van Agtmaal et al., 2012). KLF2 expression is promoted by athero-protective flow patterns and induces transcriptional changes in hundreds of endothelial genes, resulting in anti-inflammatory and antithrombotic cellular adaptations (Dekker et al., 2006; Fledderus et al., 2007; Kumar et al., 2005; Lin et al., 2005). Treatment with statins also up-regulates KLF2 expression; and their anti-inflammatory, anti-coagulant and antithrombotic effects are believed to be mediated by this transcription factor (Parmar et al., 2005; Sen-Banerjee et al., 2005). However, WPB size reduction induced by statin treatment does not require KLF2 (Ferraro et al., 2016). Altogether, a significant body of experimental evidence indicates that the size of WPBs is subject to regulation and represents a target for pharmacological intervention in haemostatic function and thrombotic risk.

We therefore screened human-licenced drugs with the aim of identifying WPB size-reducing molecules that could be rapidly repurposed as antithrombotics. We found fifty-eight drugs with this activity, the majority of which can be grouped into pharmacological classes. Some of these classes, such as microtubule depolymerizing agents and statins, have been identified by previous work (Ferraro et al., 2014; Ferraro et al., 2016). Others might be expected, due to their effects on the Golgi ribbon, as in the case of HDAC and Topoisomerase inhibitors (Farber-Katz et al., 2014; Gendarme et al., 2017). Our screen also identified compounds with pharmacology previously unknown to affect WPB biogenesis and size. Together, these findings suggest that several cellular pathways can modulate the size of WPBs produced by endothelial cells.

Multidrug resistance protein 1 (MRP1), is an organic anion transporter. Its up-regulation is responsible for the development of tumor resistance to chemotherapy, hence its name. While this activity towards xenobiotics was the first to be identified, it has become clear that MRP1 is also involved in the cellular efflux of endogenous molecules, mediating pro-inflammatory signalling pathways and may act as an oxidative stress sensor (Cole, 2014). Interestingly, twenty-two compounds, with varied mechanisms of action (**Table 1 and Supplemental Table 1**), have also been described as MRP1 inhibitors. This suggests the possibility that, in addition to their main molecular target, these drugs could affect the efflux of endogenous MRP1 signalling substrates, which regulate WPB size; a mechanism worth future investigation.

Since the screen endpoint was 24 h, the drugs listed are relatively fast-acting. Except for statins (Ferraro et al., 2016) and cardiac glycosides (see above), drug activity was documented at the single concentration used in our screen (10 µM). While such concentrations are unlikely to be used for patient administration, it is worth noting that sub-micromolar concentration treatment with simvastatin for 24 h does reduces WPB size with dramatic effects on both platelet recruitment and plasma VWF adhesion to the stimulated endothelium (Ferraro et al., 2016). It therefore cannot be ruled out that several of the drugs identified by the screen would maintain WPB size-reducing activity at lower concentrations, compatible with their use in the clinic. Statins rapidly produce anti-inflammatory and anticoagulant effects on the endothelium (Greenwood and Mason, 2007) and their acute administration in the context of percutaneous angioplasty greatly reduces post-operative myocardial infarctions (Leoncini et al., 2013). The compounding of these fast-acting effects, WPB size-reduction included, suggests that statins may represent a promising tool for acute, emergency treatment in endotheliopathies involving inflammation, coagulation and thrombosis.

As a class, the most potent WPB-shortening drugs identified in the screen, active also at sub-micromolar concentrations (**Figure 2B and 3B**), were the cardiac glycosides and cardenolides. With the caution due to their known dose-dependent cytotoxicity (Kanji and MacLean, 2012), cardiac glycosides may therefore be worth exploring in acute and chronic antithrombotic therapies.

Further to the potential toxicity associated with administration of drugs at high concentrations, we note that in vitro combination of WPB-size reducing treatments, acting through different mechanisms, can display synergy in the abatement of plasma VWF recruitment to the endothelial surface (**Supplemental Figure 1**). In a clinical setting, WPB shortening and its consequent antithrombotic effects might therefore be achieved by administering combinations of drugs at lower, non-toxic concentrations.

In conclusion, here, we report a set of licenced drugs with potential antithrombotic activity, via a novel mechanism: the reduction of WPB size and its consequence in terms of reduced adhesion of platelets and circulating plasma VWF to the UL-VWF they release.

## Author contributions

F.F., R.K. and D.F.C designed study. F.F. and J.C. did experiments. J.K-V. and F.F. analysed data. R.K. provided fundamental reagents and instrumentation. F.F and D.F.C. wrote manuscript.

## Acknowledgements

The authors wish to thank Francesca Patella, John Greenwood, Marie Ann Scully, Laura Benjamin and Martin Raff for their valuable comments on the manuscript. F. F. and D.F.C. were funded by the MRC grant MC_UU_12018/2 awarded to DFC. R.K. J.C, and J K-V were funded by the MRC grant MC_U12266B.

## Materials and Methods

### Cells

Human umbilical vein endothelial cells (HUVECs) were obtained commercially from PromoCell or Lonza. Cells were from pooled donors of both sexes expanded in our lab and used at low passage (3 to 4), within 15 population doublings since isolation from umbilical cord. Cells were maintained in HGM (HUVEC Growth Medium) with the following composition: M199 (Gibco, Life Technologies), 20% Fetal Bovine Serum, (Labtech), 30 µg/mL endothelial cell growth supplement from bovine neural tissue and 10 U/mL Heparin (both from Sigma-Aldrich). Cells were cultured at 37 °C, 5% CO_2_, in humidified incubators.

### Reagents

The library compounds (1280 FDA-approved drugs), from Prestwick Chemical, were stored at – 80 °C as 10 mM stock solutions in DMSO. For the screen, compounds were transferred to Echo Qualified 384-Well Low Dead Volume Microplates using the Echo® 520 acoustic dispenser (both from Labcyte). Antibodies used in this study were: rabbit polyclonal anti-VWF pro-peptide region (Hewlett et al., 2011), kindly provided by Dr. Carter (St. George’s University, London); mouse monoclonal anti-GM130 (clone 35) from BD Biosciences; a rabbit polyclonal anti-VWF from DAKO (cat. no. A0082). DMSO (Hybri-Max™, cat. No. D2650) and Nocodazole (cat. No. M1404) were from Sigma-Aldrich. siRNA sequences targeting Luciferase and VWF were custom synthesised and previously described(Ferraro et al., 2016). Normal pooled human plasma (Cryochek™, cat. No. CCN-15) was from Precision Biologic.

### Drug screen

HUVECs were seeded on gelatin-coated 96-well plates (Nunclon surface©, NUNC) at 15000 cells/well and cultured. After 24 h, cells were rinsed with fresh medium and then library compounds were added with the Echo® 520 (Labcyte) acoustic dispenser and medium volume per well was adjusted with a MultiFlo FX dispenser (BioTek Instruments Inc.) to 100 µL for a final compound concentration of 10 µM. Plates were prepared in duplicate. Each plate contained one column treated with negative DMSO control (0.1%, final concentration) and one column treated with Nocodazole positive control (3.3 µM, final concentration). All cells (DMSO-, Nocodazole- and compound-treated) received the same amount of DMSO vehicle (0.1% vol:vol). After 24 h treatment, cells were rinsed twice with warm, fresh HGM and fixed by incubation with 4% formaldehyde in PBS (10 min, RT).

### Dose response experiments

HUVECs were seeded in 96-well plates and pre-processed as described for the screen. Cardiac glycosides were added to cells using the Echo® 520 acoustic dispenser (Labcyte) and cells were treated as described above. Concentration of each compound ranged from 78 nM to 10 µM in 2-fold increments across the wells of the plate column. Duplicate plates were prepared. Each plate contained DMSO and Nocodozale controls as detailed for the screen. Treatment was for 24 h, at the end of which cells were fixed as described for the screen.

### Immunostaining and image acquisition

Fixed cells were processed for immunostaining as previously described (Ferraro et al., 2016). WPBs were labelled using a rabbit polyclonal antibody to the VWF pro-peptide region. The Golgi apparatus was labeled with an anti-GM130 mAb. Primary antibodies were detected with Alexa Fluor dye-conjugated antibodies (Life Technologies). Nuclei were counterstained with 33342 (Life Technologies). Images (9 fields of view per well) were acquired with an Opera High Content Screening System (Perkin Elmer) using a 40x air objective (NA 0.6).

### High-throughput morphometry (HTM) workflow

Image processing and WPB extraction of morphological parameters (high-throughput morphometry, HTM) have been described in detail elsewhere (Ferraro et al., 2014). WPB size was expressed per well (i.e., summing the values measured in the 9 fields of view) as the fraction of the total WPB area covered by WPBs > 2 µm. Dose-response data analysis was done in R language, using the DRC package by Christian Ritz and Jens C. Strebig (https://CRAN.R-project.org/package=drc). The drm () function was used to fit a dose-response model, a four-parameter log-logistic function (LL.4), applied to each dataset; four parameter values were calculated: slope, lower limit, upper limit and EC_50_ value.

### Plasma VWF recruitment assays

siRNA nucleofections, drug treatment and human plasma perfusion experiments on HUVEC monolayers were carried out as previously described (Ferraro et al., 2016).

## Supplemental material

**Supplemental Table 1.**
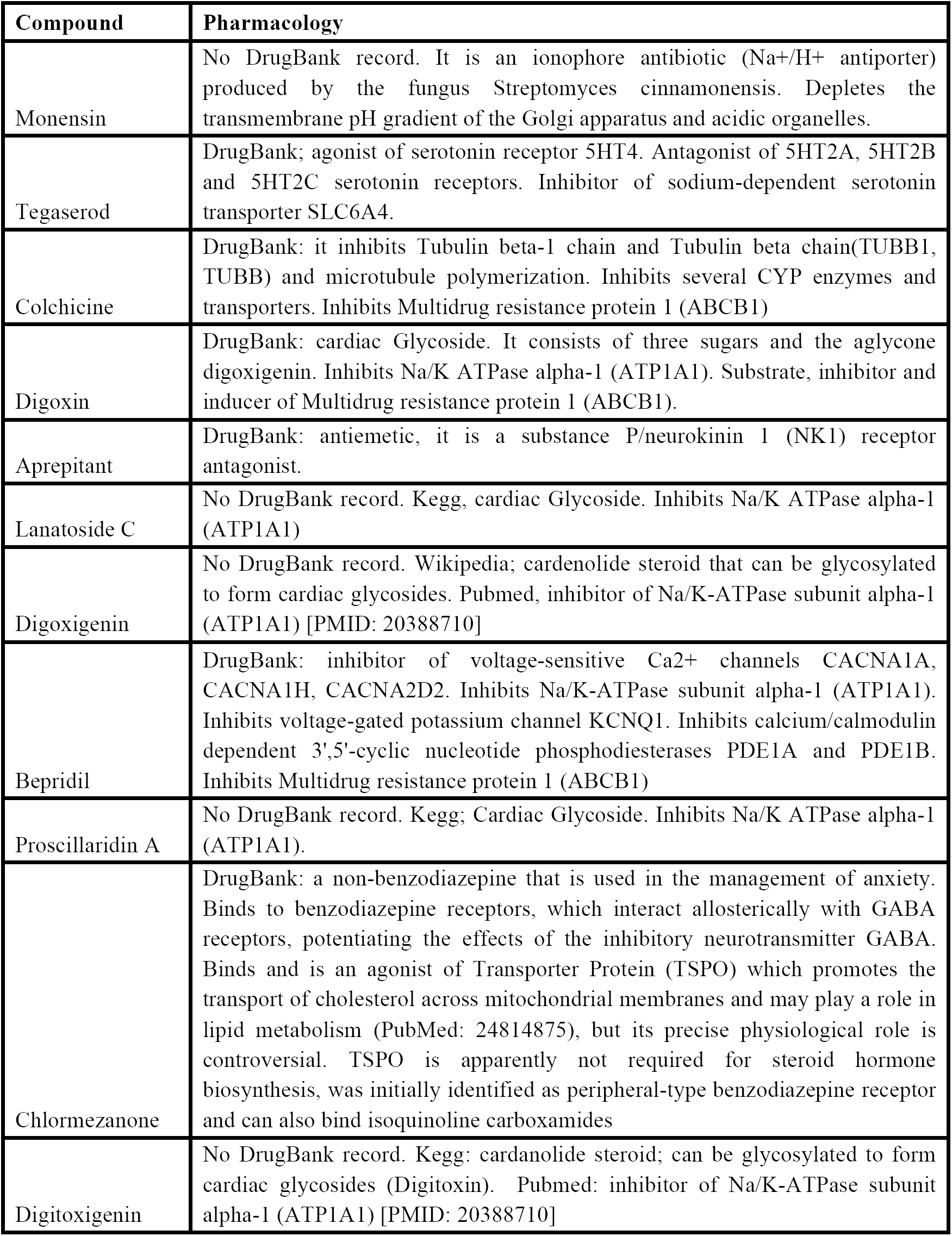
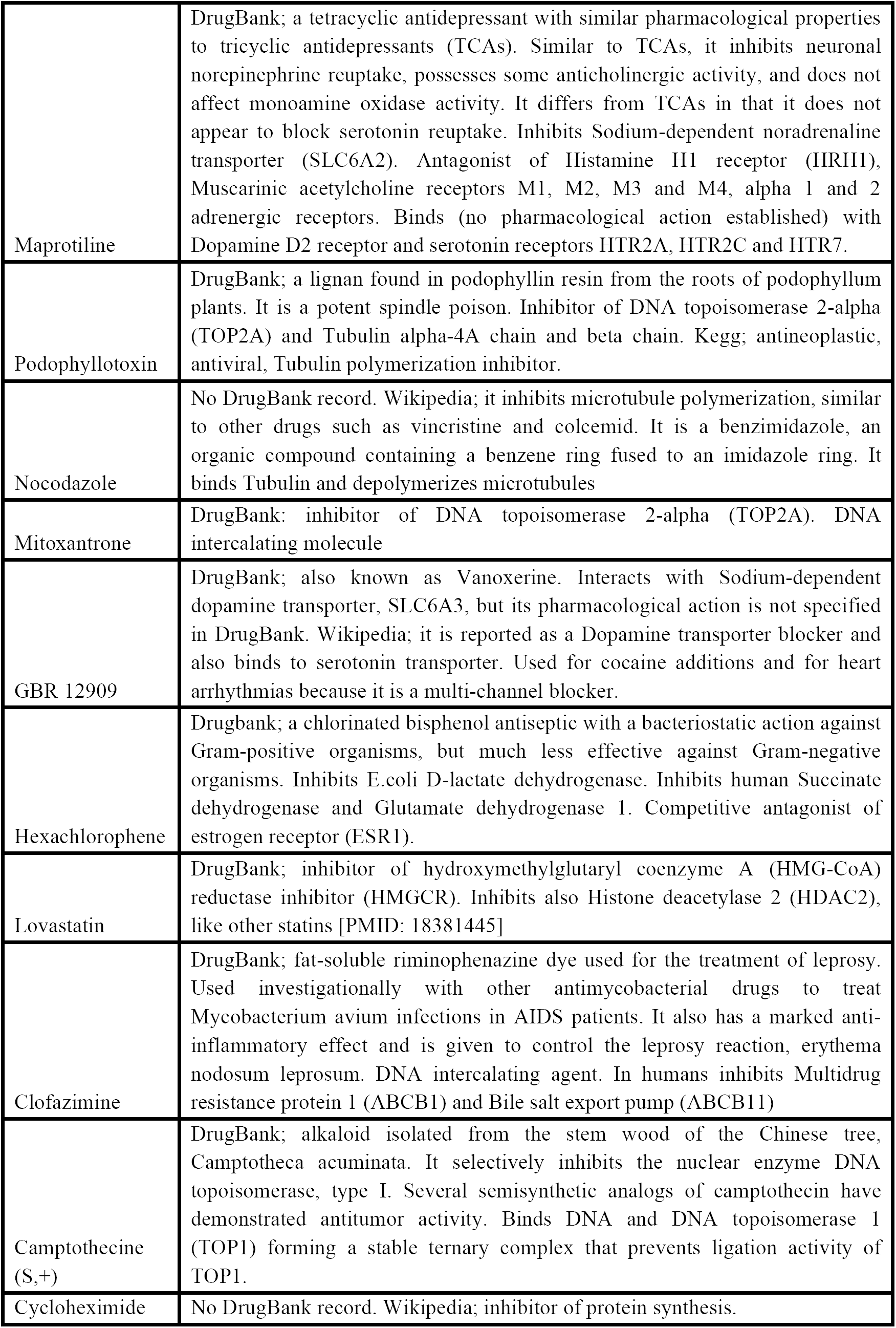
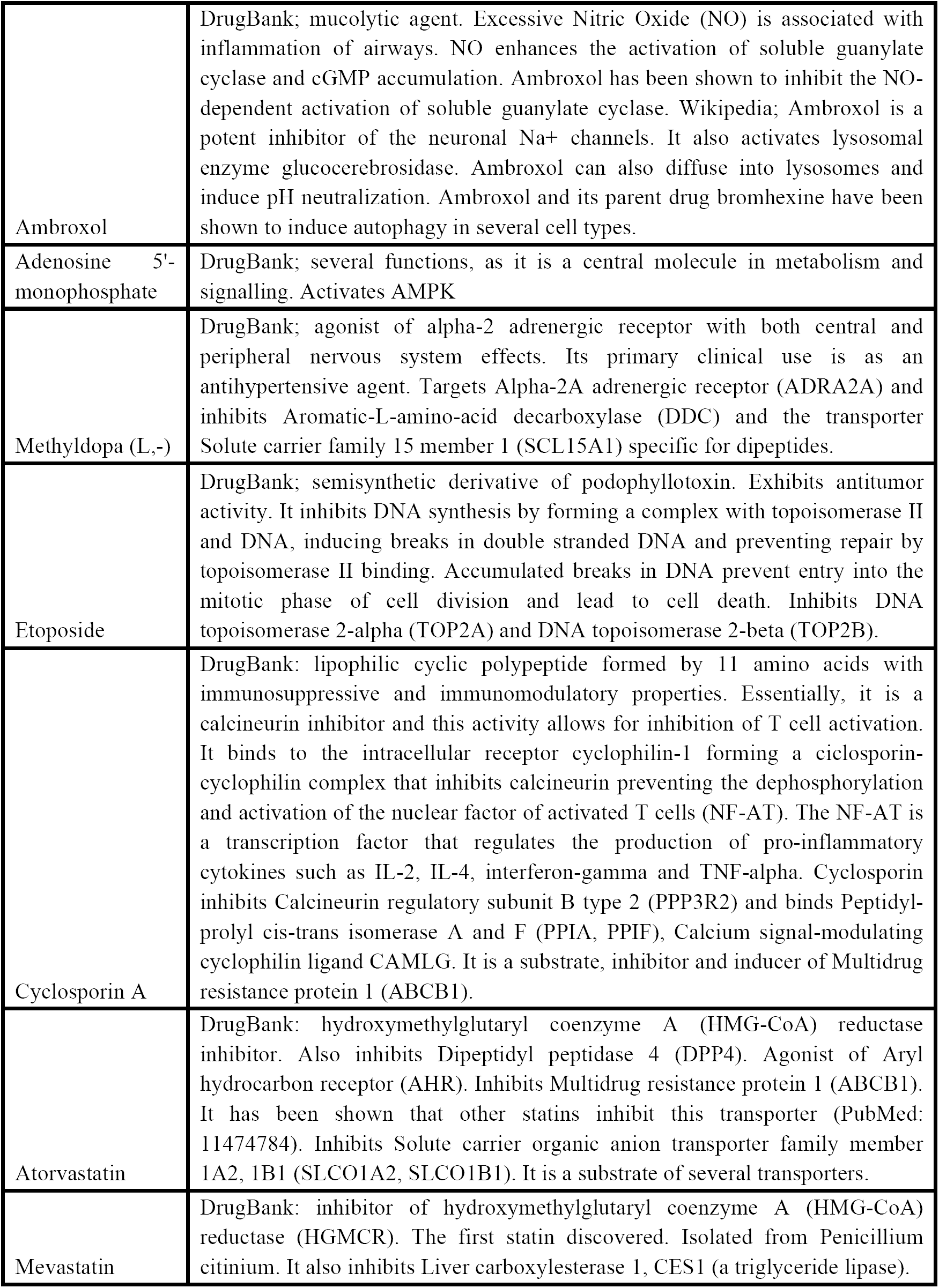
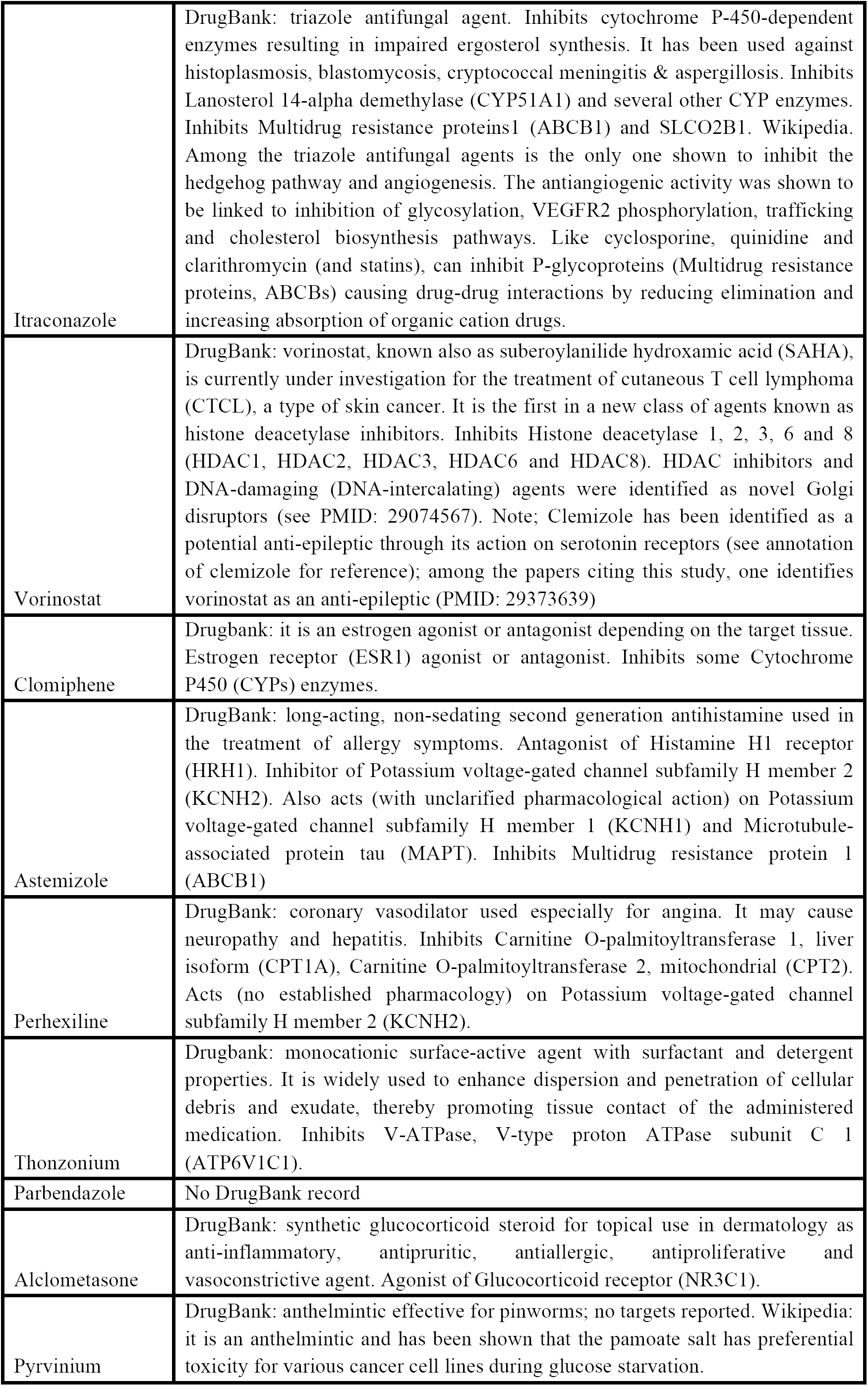
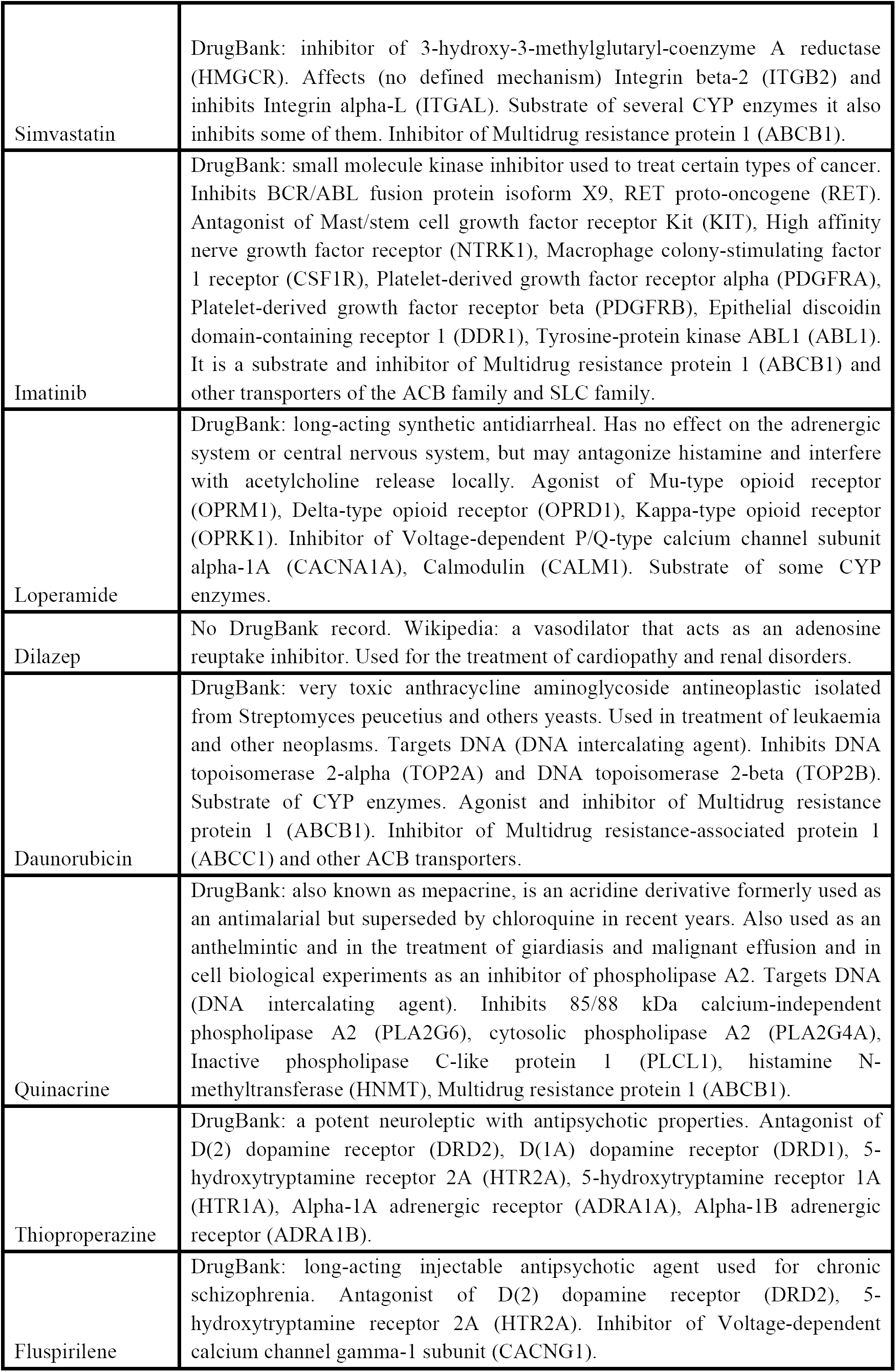
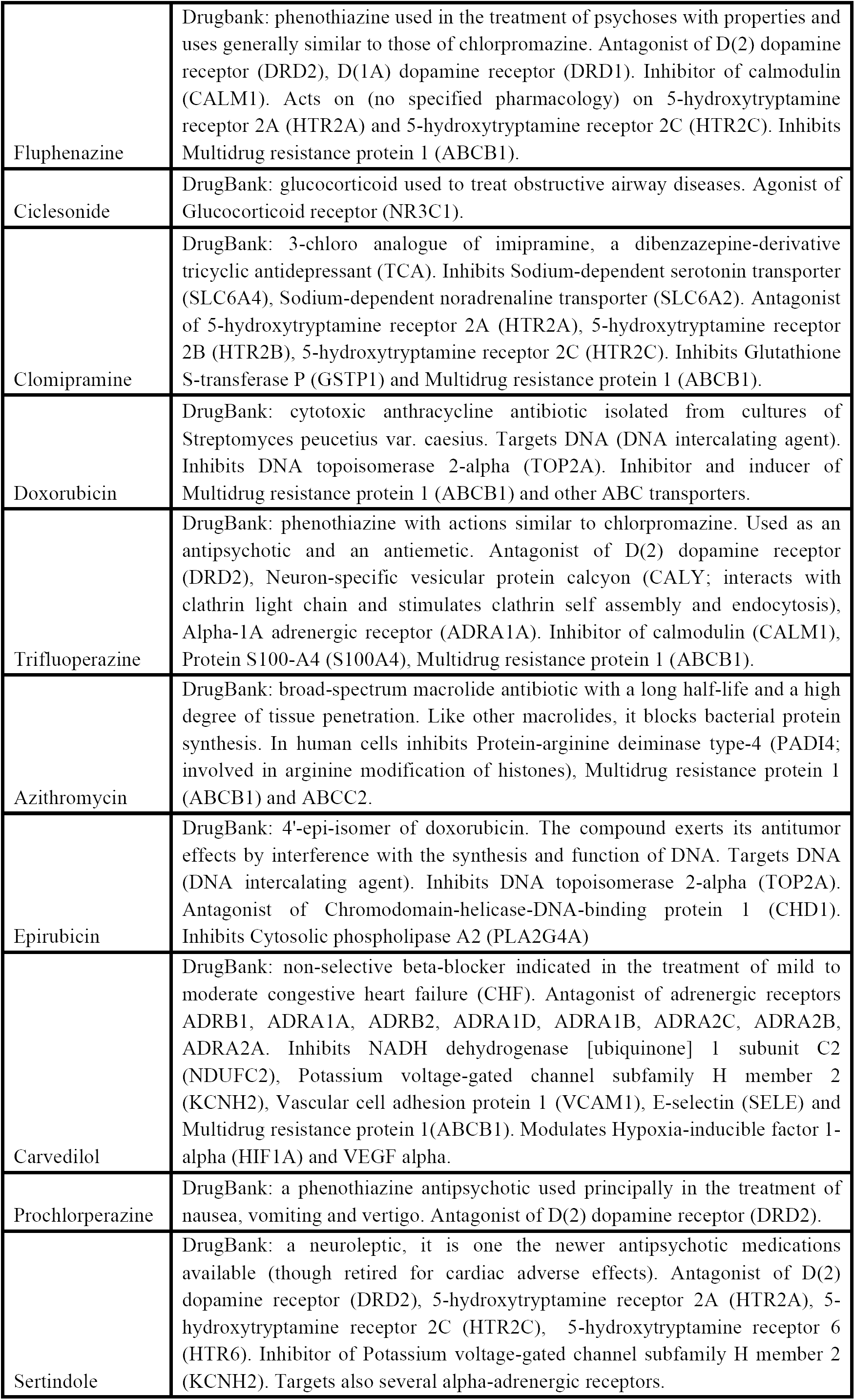
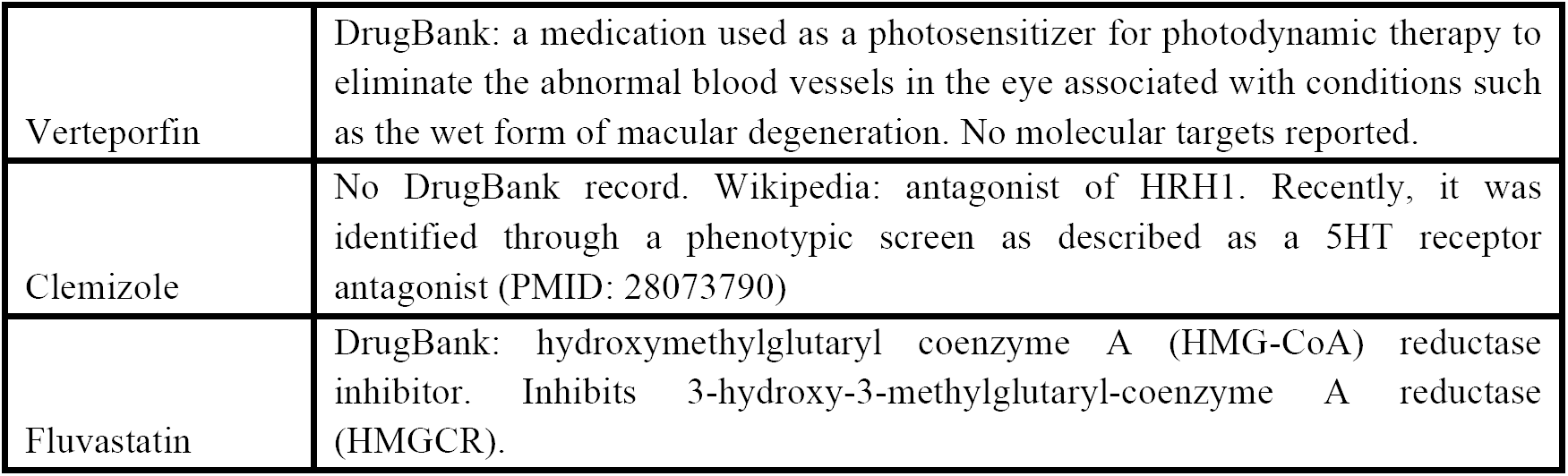
Pharmacology of hit compounds. Information regarding the mechanism of action of hit drugs was searched in PubMed and DrugBank (https://www.drugbank.ca/). Forty-one compounds can be assigned to ten pharmacological classes, each containing at least 3 drugs (see Table 1).

**Supplemental Figure 1.**
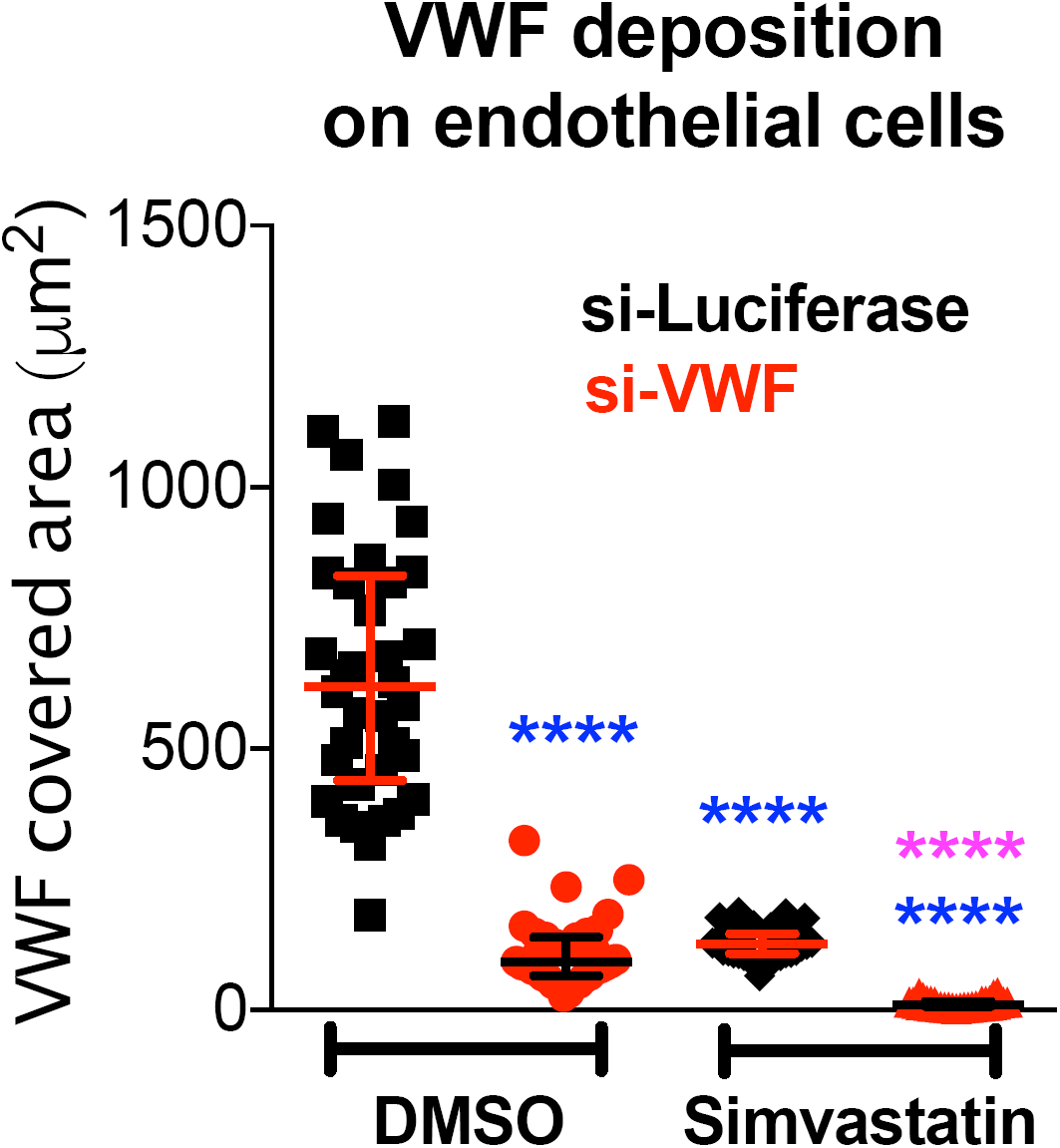
Synergistic effects of WPB-shortening treatments on plasma VWF adhesion. WPB size was reduced by two treatments, simvastatin incubation and reduced VWF synthesis (Ferraro et al., 2014; Ferraro et al., 2016), alone or in combination. HUVECs were nucleofected with siRNAs targeting luciferase (negative control) or VWF and seeded in µ-slides VI (Ibidi). At 24 h post-nucleofection, cells were treated with DMSO or 2.5 µM Simvastatin. After 24 h drug treatment, and 48 h post-nucleofection, cells were exposed to histamine to stimulate VWF secretion, while perfused with pooled human plasma and then fixed under flow (as described in Ferraro et al., 2016). VWF on the endothelial surface was detected by immunofluorescence; the area covered by its signal measures the extent of its adhesion. Each data-point represents the quantification of a field of view; median and interquartile ranges are shown. ****, P < 0.0001, Mann-Whitney test. Blue asterisks: comparisons to si-Luciferase/DMSO treatment. Purple asterisks: comparison to si-VWF/Simvastatin treatment. One of two independent experiments with similar results is shown.

